# On the Stability Landscape of the Human Gut Microbiome: Implications for Microbiome-based Therapies

**DOI:** 10.1101/176941

**Authors:** Travis E. Gibson, Vincent Carey, Amir Bashan, Elizabeth L. Hohmann, Scott T. Weiss, Yang-Yu Liu

## Abstract

Understanding how gut microbial species determine their abundances is crucial in developing any microbiome-based therapy. Towards that end, we show that the compositions of our gut microbiota have characteristic and attractive steady states, and hence respond to perturbations in predictable ways. This is achieved by developing a new method to analyze the stability landscape of the human gut microbiome. In order to illustrate the efficacy of our method and its ecological interpretation in terms of asymptotic stability, this novel method is applied to various human cohorts, including large cross-sectional studies, long longitudinal studies with frequent sampling, and perturbation studies via fecal microbiota transplantation, antibiotic and probiotic treatments. These findings will facilitate future ecological modeling efforts in human microbiome research. Moreover, the method allows for the prediction of the compositional shift of the gut microbiome during the fecal microbiota transplantation process. This result holds promise for translational applications, such as, personalized donor selection when performing fecal microbiota transplantations.

**One Sentence Summary:** A new method for analyzing the stability landscape of the human gut microbiome and predicting its steady-state composition is developed.

## 1. Introduction

Thanks to recent advances in high-throughput sequencing technologies, we now have a good understanding of what microbes reside in, and on, our bodies and their relative abundances (1, 2). Longitudinal studies have shown that the gut microbiome compositions of healthy adults are fairly consistent over time (3–5). In addition, the gut microbiome compositions of the same individual at different time points are more similar to one another than to the gut microbiome compositions of different individuals (4). This temporal stability suggests the existence of personalized stable steady states for the human gut microbiome. The landscape of stable states for the human gut microbiome is still unknown, however (39,49). It remains unclear if the gut microbiome of a healthy adult is just *marginally stable* (Fig.1a) or, if in addition, it is *asymptotically stable*, i.e., displays the feature of attractivity so that stable states behave as attractors (Fig.1b). In other words, we want to know if the landscape of stable states is *flat* or *concave*. We are also interested in knowing whether true multi-stability exists for the human gut microbiome (if there are multiple attractors for any given collection of species).

**Fig. 1.**
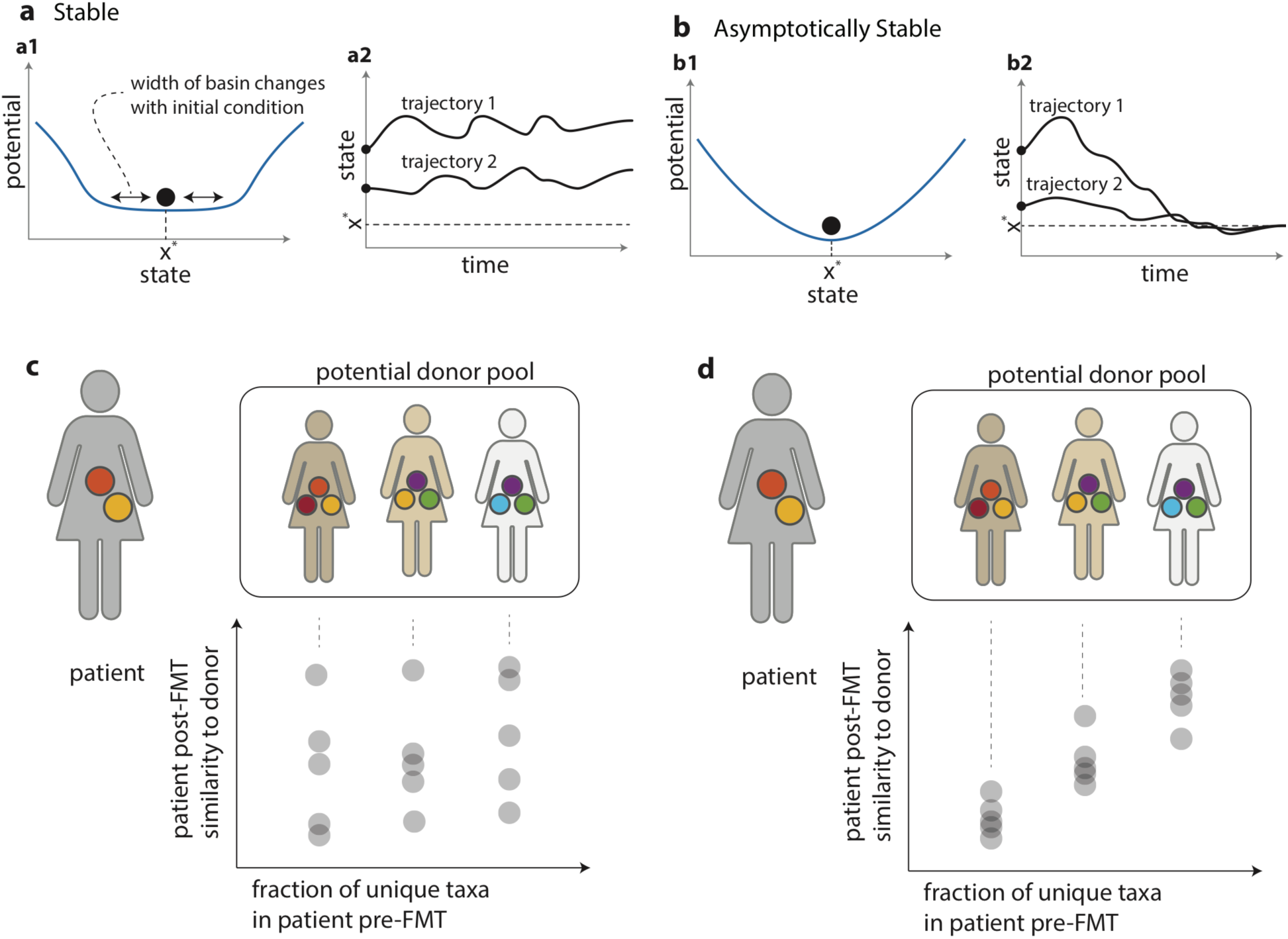
Stability schematic and FMT donor selection schematic. (a) Illustrations of the definition of stability in terms of (a1) a stability landscape (45) and (a2) trajectories near steady state. Note the lack of attractivity to the steady state. (b) Illustrations of the definition of asymptotic stability in terms of (b1) a stability landscape (45) and (b2) trajectories near steady state. Notice that even with different initial conditions the steady state is reached by both trajectories. (c) An illustration of our ability to predict the post-FMT steady state from pre-FMT analysis of samples if the dynamics are merely stable. Similarity in collection need not imply similarity in abundance profile post FMT. (d) An illustration of our ability to predict the post-FMT steady state from pre-FMT analysis of samples if the dynamics are asymptotically stable. The more similar the patients gut is in terms of taxa to the donor, the more similar the post-FMT abundance profile will be to the donor.

Addressing these fundamental stability-related questions will help in deciphering the complex microbial dynamics underlying the human gut microbiome and will facilitate future ecological modeling efforts. Moreover, it will enable us to predict the steady-state composition of human gut microbiota based on species collection alone. This has immediate translational implications. Consider for instance the case of *Clostridium difficile* infection (CDI), which accounts for 15 to 25% of antibiotic-associated diarrhea, and is an increasing health problem in the United States, leading to nearly 500,000 diagnoses and approximately 30,000 deaths every year (40). In a healthy gut microbiota, *C. difficile* is out-competed by hundreds of strains of bacteria that are normally present. However, after repeated broad-spectrum antibiotic administrations that disrupt the healthy microbial community, *C. difficile* can flourish within the gut and produce potent toxins. Because *C. difficile* can form spores that are not killed by antibiotics, and because the normal microbiota is diminished, the infection is often poorly or transiently responsive to standard antibiotics such as oral vancomycin or metronidazole and recurs. In contrast, after hundreds of treatments in many independent institutions, fecal microbiota transplantation (FMT) has been shown to cure over 90% of the most recalcitrant CDI cases that had previously failed standard antibiotic therapy (41). A schematic of microbial abundances throughout the FMT process is shown in Supplemental Figure S1. Despite these promising findings, long-term safety concerns still linger and there is still no clear criterion for personalized donor selection when given a pool of healthy donor candidates. Indeed, rather simple questions remain unanswered. For example, it is unknown whether the patient’s gut microbiota return to an abundance profile similar to his/her pre-antibiotic state, or become more similar to that of the donor’s. The work presented here will help us predict the compositional shift of the patient’s gut microbiome during FMT, and hence facilitate personalized donor selection.

Stability is a fundamental notion in dynamical systems theory. For a general dynamical system of *n* components, the temporal behavior of the system’s *n*-dimensional state vector ***x*** is usually described by a set of ordinary differential equations d***x***/d*t* = ***f*** (***x***, *t*), where ***f*** (***x***, *t*) determines the “rules” governing the temporal behavior of this system. In the context of population dynamics, the state vector ***x*** represents the abundance profile of the present species, and ***f*** (***x***, *t*) may capture the intrinsic growth of the species and the intra- and inter-species interactions. The mathematical foundations of stability were given by Lyapunov (7–11). A variety of stability-types exist for a general dynamical system. Roughly speaking, if all solutions of the equations d***x***/d*t* = ***f*** (***x***, *t*) that start out near an equilibrium point ***x***\* (which satisfies ***f*** (***x***\*, *t*)=0) stay near ***x***\* forever, then ***x***\* is *stable*. More strongly, if ***x***\* is stable and all solutions that start out near ***x***\* converge to ***x***\* as *t* → ∞, then ***x***\* is *asymptotically stable*. ***x***\* is *marginally stable* if it is stable but not asymptotically stable. The key difference between marginal stability and asymptotic stability is that the latter has an *attractivity* feature such that the equilibrium point ***x***\* is “attracting” nearby states while the former doesn’t have this feature, see Figure 1c and 1d, as well as the methods section for more detailed definitions and a discussion about perturbed and unperturbed dynamics, along with reference note (55).

A straightforward approach to studying the stability landscape of the human gut microbiome would be to infer and study the underlying ecological dynamics. Yet, this is very challenging, if not impossible, given the issues surrounding compositionality of relative abundance data and the lack of sufficient variation in existing temporal metagenomics data (6, 28, 29). Collecting temporal data with more variation requires well-designed perturbation experiments (56), which may raise serious ethical concerns. In this work a new method to characterize the stability landscape of the human gut microbiome without inferring the underlying ecological dynamics is developed. In particular, through the development of appropriate dissimilarity measures, we show that the human gut microbiome actually demonstrates not only stability but also the feature of attractivity. In the following sections the attractivity feature of human gut microbiota is demonstrated through the visualization of longitudinal data. Asymptotic stability is then verified by applying the developed dissimilarity measures to large cross-sectional cohorts (55). Finally, we demonstrate our ability to predict patients’ post-FMT microbiota similarity to that of their donors by analyzing patients’ pre-FMT stool samples alone, paving the way for personalized donor selection in FMT.

## 2. Method

### 2.1. Demonstration of asymptotic stability in longitudinal microbiome data

The notion of stability in the context of population dynamics and ecological systems has been discussed since 1839 (12–15) and are ongoing (16–19,42,51-54). Stability’s importance in the context of the human gut microbiome has been recently recognized as well (20–22, 50). Indeed, there can be day-to-day perturbations that drive an individual’s abundance profiles away from the unperturbed steady state (25). However, upon closer inspection of the state trajectories in (25, Figure 3A) it was found that the trajectories always returned to the same steady region (26). To further investigate this attractive feature of the human gut microbiome, the nearly daily abundance profiles from male and female subjects in the Moving Pictures Study (MPS) (23) were projected onto the principle component space of more than 6,000 samples from the American Gut Project (AGP) (27), see Figure 2 and Supplemental Movies 1 and 2. We find that over a typical 10 to 20-day window, the abundance profile of each subject stays in the same region in the principle component space. Large deviations from that region do occur to the male subject, possibly due to external perturbations (e.g., diet change), but then the system quickly returns to the same region. This is an illustration of the attractivity feature of a dynamical system for which the unperturbed dynamics are asymptotically stable (55).

**Fig. 2.**
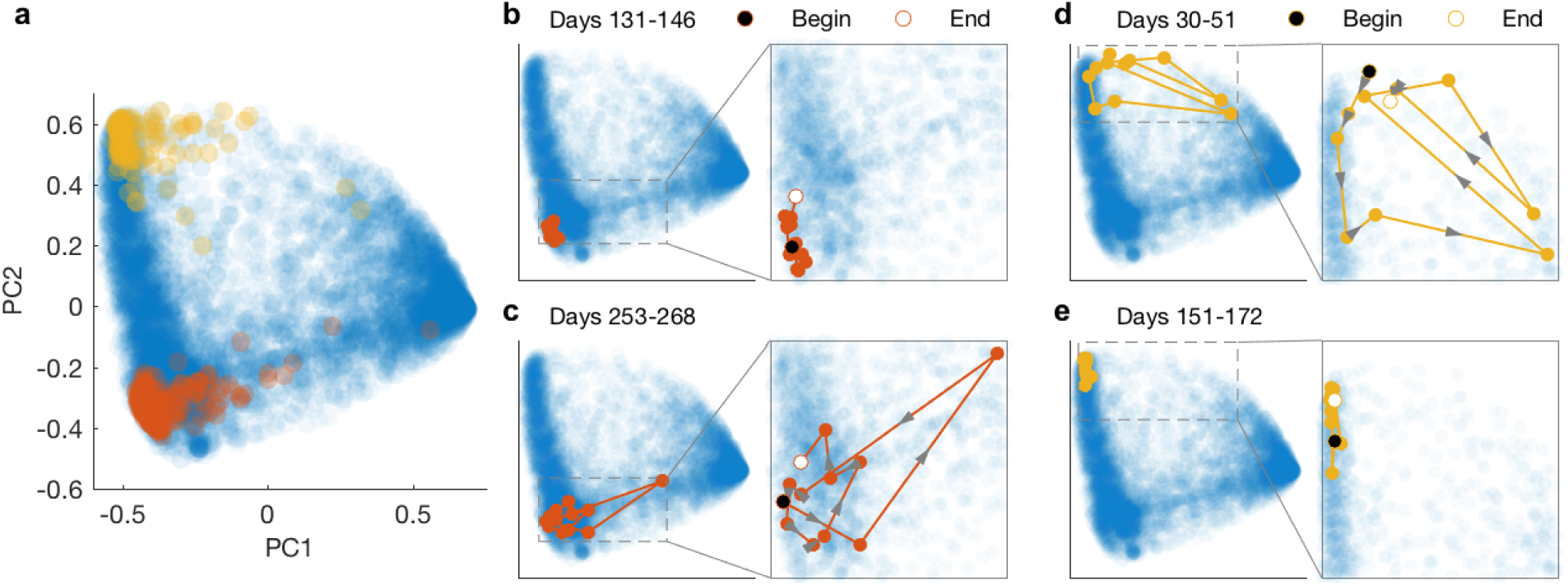
Attractivity Visualized through Longitudinal Data. (a) Background in blue is scope of variation for the American Gut Project (AGP, cross sectional cohort of n = 6,375). Orange Dots are from a single male subject, (n = 336 longitudinal almost daily samples), and yellow dots are from a female subject, (n = 131 longitudinal less frequently sampled but over the same time window) and are from the Moving Pictures Study (MPS). AGP and MPS samples were analyzed using the same OTU picking scheme, principal components were derived from the AGP samples only and then the male and female longitudinal samples were projected into PC1 and PC2. (b) A typical multiple day snapshot of the male samples with all samples staying in the same region. (c) An instance where the samples deviated from the steady region for the male, but then returning to the steady region rapidly after each deviation. (d) An instance where the samples deviated from the steady region for the female, but then returning to the steady region within a few days after each deviation. (e) A typical multiple day snapshot of the female samples with all samples staying in the same region.

**Fig. 3.**
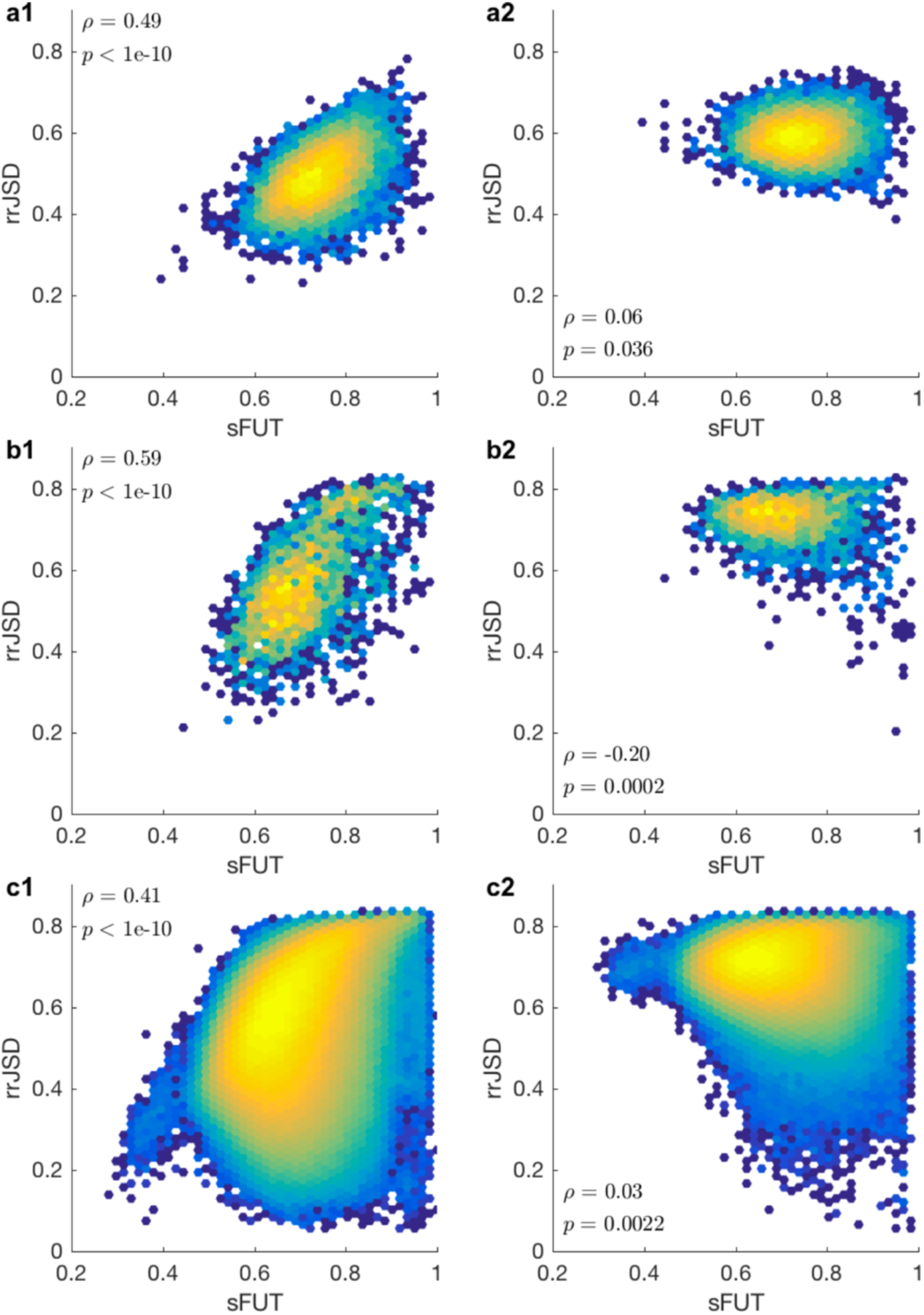
Hexagonal Density Plots for Fraction of Unique Taxa versus Similarity in Healthy Cross-sectional Cohorts. Correlation coefficient, ρ, and boot strapped p-values displayed, see Methods section for details. (a) HMP fecal samples, n = 146 resulting in 10,585 sample pairs (a1) original samples (a2) randomized samples. (b) SMP data, n = 72 resulting in 2,556 sample pairs (b1) original samples (b2) randomized samples. (c) Healthy AGP data, n = 2,597 resulting in 3,370,906 sample pairs (c1) original samples (c2) randomized samples. All healthy cohorts illustrate the strong correlation between sFUT and rrJSD showing that the less unique taxa between any two samples the more similar the abundance profiles.

### 2.2. Detection of asymptotic stability from cross-sectional microbiome data

Results from the two individuals shown in Fig. 2 leads to questions best answered by a large cohort of *longitudinal* data. However, this kind of data is simply not available at present. To resolve this issue, we now introduce a new method that can be used to analyze *cross-sectional* microbiome samples. The method characterizes how species collection alone affects an individual’s microbiota composition. To achieve this, two classes of dissimilarity measures are introduced for comparing microbiome samples.

The first class of dissimilarity measures quantify the Fraction of Unique Taxa (FUT) between two samples. This is similar to the classic Jaccard distance, however this method uses a different denominator for normalization, because the normalization used in Jaccard is biased towards large sets (i.e., samples with more present OTUs), see the Methods Section. We use two functions to quantify FUT: sFUT(*x*, *y*) and aFUT(*x*, *y*). The distance metric sFUT(*x*, *y*) is symmetric and is simply the total number of unique taxa between two samples *x*, *y* normalized by the total number of taxa in both samples (note that this is not the union). The quasi-metric aFUT(*x*, *y*) is asymmetric and is defined as the total number of unique taxa in *x* relative to *y*, normalized by the average number of taxa in the pair (*x*, *y*). For rigorous definitions see the Methods Section. The dissimilarity measure aFUT(*x*, *y*) is designed specifically for the analysis of pre-FMT patient and donor stool samples. In that case, we only account for the unique taxa in *x* (the patient’s pre-FMT sample), because after FMT the taxa within the donor that were unique have now been introduced to the patient. With the use of aFUT(*x*, *y*) one can predict post-FMT similarity between donor and patient from pre-FMT fecal sample analysis alone, offering a simple practical tool for donor selection.

The second class of dissimilarity measures is motivated by our renormalization technique applied in (6), where spurious correlations due to the compositionality of genomic survey data are removed. Given a sample pair (*x*, *y*) we first exclude all taxa not shared by the samples *x* and *y*. Then we renormalize the abundances of the shared taxa and quantify the dissimilarity between two renormalized abundance profiles using any classical measures, e.g., Euclidean, Jensen-Shannon Divergence (JSD), root of JSD (rJSD), Bray-Curtis, Yue-Clayton, etc.. (In this work, rJSD is used, simply because it is a distance metric.) The important step here is the *renormalization* procedure. To emphasize this point, we use rrJSD to denote the rJSD over renormalized abundance profiles. For more details on rrJSD see the Methods Section.

Using the above two measures it is then possible to assess the relationship between collection and abundance. If collection is a dominant factor in determining abundance, then there will be a strong degree of correlation between sFUT and rrJSD. Interpreting this relationship then allows one to make claims regarding the stability characteristics of the human gut microbiome, as is done in the following section.

## 3. Results

### 3.1. Healthy human gut microbiota

In Fig. 2 we showed how the longitudinal data of two human subjects qualitatively supports the notion of asymptotic stability. Here we try to add more quantitative support to this claim by analyzing cross-sectional data from large healthy cohorts of subjects. After applying sFUT and rrJSD distances to all sample pairs in three large cross-sectional cohorts (The Human Microbiome Project [HMP], the Student Microbiome Project [SMP], and the AGP), we observe a strong correlation between sFUT and rrJSD as illustrated by Figure 3, and Supplemental Figure S3. This strong correlation suggests that the more similar two samples are in terms of taxa, the more similar their renormalized abundance profiles are. For a system that is asymptotically stable we would expect this strong correlation, but for a marginally stable system that does not have this attractivity feature this trend is not expected, see Supplemental Figure S4.

These results also suggest that for each collection of taxa in the human gut microbiome there is a unique stable state (that is, in the absence of perturbations there is a unique fixed point, but what we observe is a constantly perturbed system near the unperturbed fixed point). If each fixed collection could support multiple stable states (i.e., true multi-stability exists), then sFUT and rrJSD would not be strongly correlated. Indeed, alternative stable states for the same collection of taxa would behave as different attractors of the microbial dynamics, and present as a multimodal distribution for low values of sFUT. Those samples taken from the same attractor would be very similar in rrJSD and those samples taken between two attractors would be less similar in rrJSD. Note that after altering the collection of microbes, for instance through antibiotic or probiotic administration (24), the microbiome can settle into a new steady abundance profile. This does not imply the existence of true multi-stability or alternative stable states, however. When new species are introduced into an ecosystem, the dimension of the state space increases, see Supplemental Figure S2. The higher dimensional system could have a unique stable state, which will look like an alternative stable state in the lower dimensional projected state space. Therefore, it is important to take species collection into account when discussing the stability landscape of the human microbiome, a factor that has not always been accounted for, even though it is fundamental to the issue.

### 3.2. Perturbed Human Gut Microbiota

The large cross-sectional studies provide indirect yet quantitative evidence for the asymptotic stability of the gut microbiome of healthy adults. In order for this result to be useful, and to further illustrate evidence for asymptotic stability in the human gut microbiome, the strong correlation seen in the healthy cross-sectional cohorts should also hold when perturbations are applied directly to the collection of taxa. One important feature of an asymptotically stable ecosystem is robustness in terms of initial abundances and convergence to the steady state of the system. In other words, the specific initial abundances of the present taxa will not be as important as the actual presence or absence of the taxa in the community, see Figure 1b. In the following sections we will continue to illustrate the asymptotic stability of human gut microbiota by analyzing perturbed microbiomes. The perturbations analyzed are: antibiotic administration, probiotic supplementation, and FMT. Regarding FMT, we are also able to predict the similarity of a patient’s post-FMT microbiome to that of the donor by analyzing the patient’s pre-FMT fecal samples.

#### 3.2.1. Perturbation by Antibiotics

The distance functions are now applied to a longitudinal cohort of subjects who received antibiotics. In (24) three patients were followed for 10 months while antibiotics (ciprofloxacin) were administered at the two-month time point and again shortly after the beginning of the eighth month. Our concern is the change in steady state of those individuals, and thus we are only interested in the three stable phases: the time period prior to the first dose of antibiotics (phase 1), the time period one week after the end of the first round of antibiotics, up until the second round of antibiotics was administered (phase 2), and then the time period starting one week after the last round of antibiotics ended, up until the end of the experiment (phase 3). The three phases correlate to Pre Cp, Interim, and Post Cp respectively in the original study (24). Difference in collection using sFUT versus abundance dissimilarity using rrJSD between all sample pairs in the three phases described above is shown in Figure 4. In Figure 4a data for all sample pairs is shown with the intra-patient samples highlighted in blue and the inter-patient samples highlighted in orange. The intra-patient pairs are more similar in terms of taxa, which is to be expected, and consistent with what we know already: within patient variation is smaller than between patient variation (4, 23, 24). The trend that rrJSD is strongly correlated with sFUT for all sample pairs is also consistent with the cross-sectional data analysis in Figure 3. In Figure 4b, only intra-patient samples are shown, with the sample pairs highlighted by patient. The correlation between rrJSD and aFUT is thus independent of the patient. While the patient’s response to the antibiotics is indeed personalized in terms of the specific taxa lost during each round of antibiotic administration (24), the correlation between the sample pairs is preserved.

**Fig. 4.**
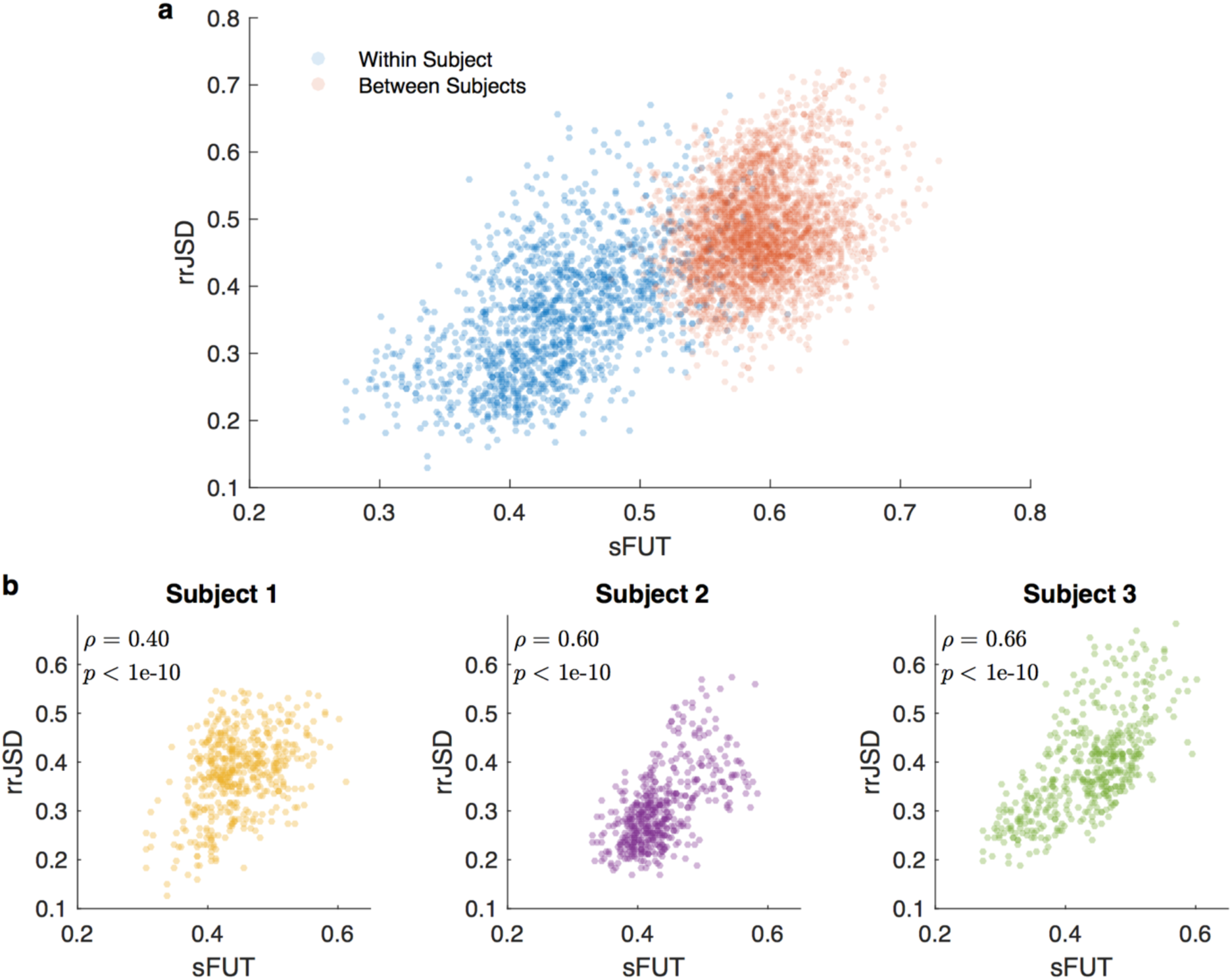
Fraction of Unique Taxa versus Similarity in Healthy Individuals taking Antibiotics. Healthy individuals were given two rounds of antibiotics each. The data shown here represents the data before the first round and at least one week after each round of antibiotics and no data is from the time period when antibiotics were taken. Thus the data is more or less a representation the three steady time periods for the individuals. The number of samples meeting this criterion is 32, 21, and 31 for the three individuals respectively. (a) sFUT versus rrJSD between subjects, highlighted in red resulting in 2315 sample pairs, and within subjects, highlighted in blue resulting in 1,171 sample pairs (b) Fraction of Unique Taxa versus Similarity for Subject 1, resulting in 496 sample pairs, Subject 2, resulting in 210 sample pairs, and Subject 3, resulting in 465 sample pairs. Correlation coefficient, ρ, and boot strapped p-values displayed, see Methods section for details. The same trend as seen in the healthy cohorts in Figure 3 is seen here as well with samples within subjects more similar than between subjects. The response from each individual to the antibiotics may be unique in terms of the specific taxa that are affected, but the overall trend in terms of the two distances is the same for each individual.

Hence, even if we are not able to predict what taxa will be removed during each round of antibiotics, knowing which taxa are present enables the prediction of a patient’s abundance profile (either from a cross-sectional cohort or from their own longitudinal samples). This process of prediction will be demonstrated with FMT samples in a later section.

#### 3.2.2. Perturbation by Probiotics

Next, we discuss two longitudinal cohorts that received probiotics (30, 31). In (30) the commercial product Enterolactis Plus was used, which contains the single strain *Lactobacillus paracasei* DG. In (31) a non-commercial probiotic was used which contained the single strain Bifidobacterium bifidum Bb. In each of the original studies there are two groups: A and B. A pre-probiotic administration fecal sample is obtained, sample v_1_, followed by 4 weeks of probiotics for group A and 4 weeks of placebo for group B. At the end of the four weeks, another fecal sample is obtained, v_2_. Then there is a 4-week wash out period, another sample is taken, v_3_, then the placebo and probiotic are switched between the groups (crossover study design) with four weeks of application, and then a final sample v_4_ is taken. For each study, clustering analysis was performed on all fecal samples and then projected into the first two principal components, see Figures 5a1 and 5b1. The labeled data in the same two-dimensional projection is shown in Figures 5a2 and 5b2. Note that the probiotic samples are equally distributed throughout the projection space and are also equally distributed between the two clusters in both studies. More interestingly, the strong correlation between rrJSD and sFUT that was observed in the cross-sectional cohorts and the antibiotics study are observed here again for the probiotics studies. This suggests that similarity in collection implies similarity in abundance for each individual throughout the experiments, see Figures 5a3 and 5b3. Note that here, instead of computing sFUT and rrJSD between all sample pairs within a patient, we compared the three different phases of the trials to the initial condition of the respective patients, so as to illustrate the way different perturbations affected the abundance profiles (correlation coefficient, ρ, was calculated using data from the three phases together). So as to ensure a Yule– Simpson effect is not occurring the confidence intervals for the independently calculated correlation coefficients are also reported in the figure, and indeed all three phases have positive correlations. This confirms in a way what was demonstrated in Figures 5a1,b1 and 5a2,b2. The probiotic phase of the study has an indistinguishable impact on the overall abundance profiles within the patients. Thus, while probiotics may have a targeted impact on the specific biochemical pathways of interest, we do not detect a global population level impact on the microbiome.

**Fig. 5.**
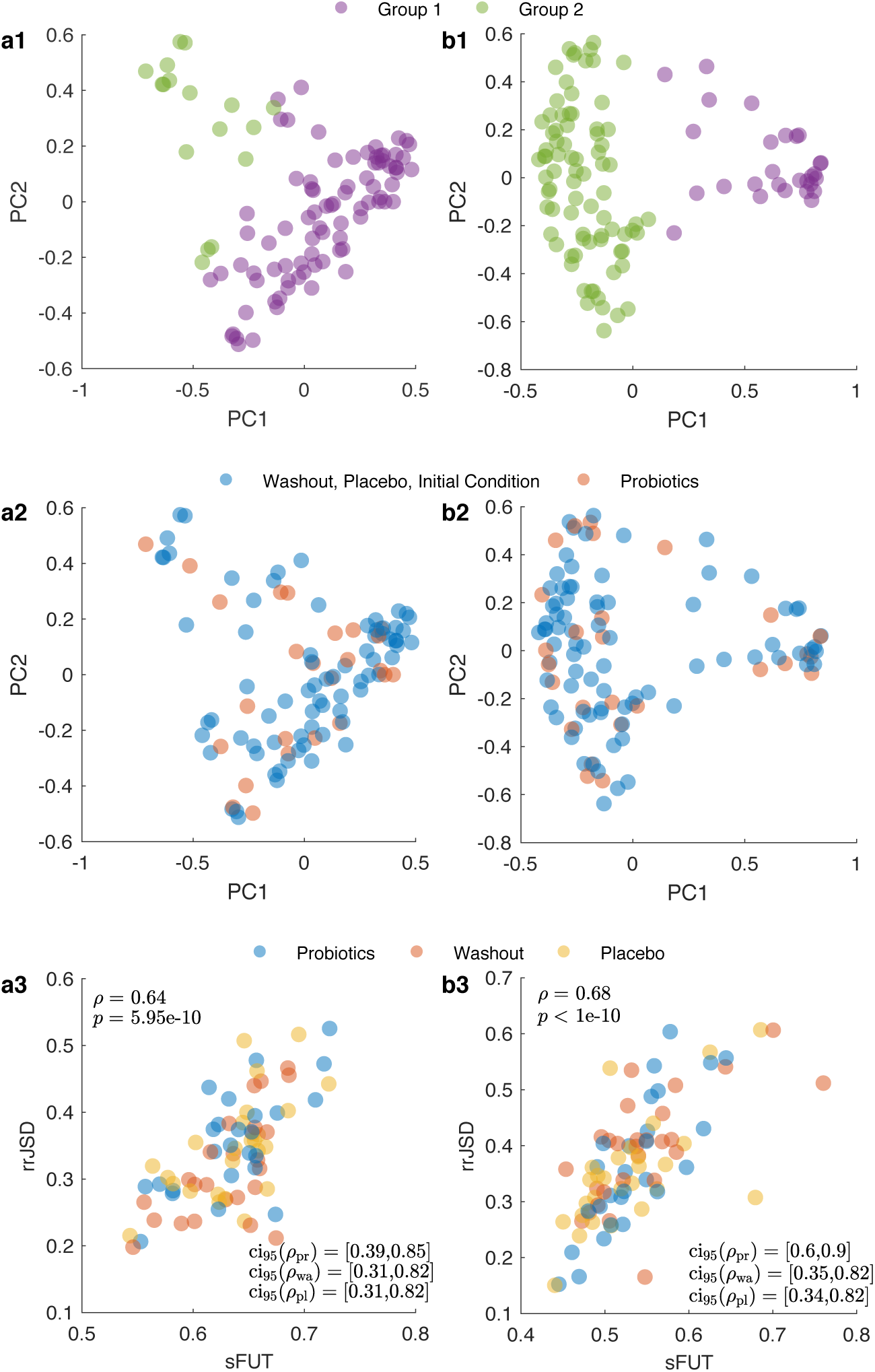
Perturbation by probiotics (a) Cross-over study from (30) for 25 patients resulting in n = 100 samples. (a1) Clustering performed by k-medoids and projected into first two principle components (a2) Labeled data with the same 2D projection. (a3) Intra-individual sFUT versus rrJSD comparing each of the visits in {v2, v3, v4} to the initial sample v1. Correlation coeficicents, ρ, combining all three phases of the trial with corresponding p-values (standard), and 95% confidence intervals for correlation within the three phases {probiotic, washout, placebo} denoted {ci_95_(ρ_pr_), ci_95_(ρ_wa_), ci_95_(ρ_pl_)}, see methods section for details. (b) Cross-over study from (31) for 27 patients resulting in n = 108 samples, with the same subfigure descriptions.

#### 3.2.3 Perturbation by FMT

FMT has been demonstrated to be a highly effective therapy for one well-known dysbiotic state, recurrent *Clostridium difficile* infection (rCDI), with an overall cure rate ∼ 90% (43). Note that the overall cure rate includes patients with subsequent FMTs if a single FMT fails. The cure rate with a single FMT ∼70%-80% (36,41). FMT has also been proposed as a treatment for other gastrointestinal (GI) diseases such as inflammatory bowel disease (IBD), irritable bowel syndrome (IBS), and chronic constipation, as well as a variety of non-GI disorders as divergent as autism and obesity (44). Currently, there are general guidelines for what makes a potential donor healthy enough to participate in an FMT (32–34). There are currently no guidelines, however, for matching an rCDI patient to a specific donor when there is a pool of healthy donors to choose from. Indeed, even with the same donor, different patients can have very different post-FMT steady-state abundance profiles (35), which suggests there may not be such a thing as a “super donor” (32). A personalized donor should be chosen, however, in order to have more control over the post-FMT steady-state abundance profile of an rCDI patient. We hypothesize that greater long term success is possible if the patient remains at a similar abundance profile to the donor. First and foremost, it might be that engraftment/restoration of flora is more successful when the patient has a lower aFUT relative to the donor, increasing the cure rate even for a single FMT. One can also hypothesize that there may be less risk of recurrence of CDI or ongoing non-specific GI symptoms, or post-infectious IBS, which are common in people with recurrent CDI. Understanding the role that the collection plays in determining microbial abundance could also eventually allow the repertoire of FMT applications to be expanded to other clinical conditions. First, one might consider and study the use of FMT in a first episode of CDI, and then extend FMT to other complex illnesses related to a disrupted microbiome such as IBD or Crohn’s disease, which do pose as greater challenges.

In this section we will predict the similarity of the post-FMT gut microbiota of an rCDI patient to their donor by analyzing their pre-FMT stool samples. To illustrate this, consider the trajectories shown in Figure 6a1 and Figure 6a2. The two trajectories were generated from data in (3) where the abundance profile of patients was followed for several months after the FMT. For both patients, the abundance profiles are like the donor immediately after the FMT (figure 6a1 and a2). While the purple trajectory remains similar to the donor, the blue trajectory settles on a steady state that is far from the donor. If we compare the post-FMT samples to the single donor samples as in Figure 6a3 then the same strong correlation between rrJSD and sFUT as shown before holds as well, which implies that similar taxa have a similar steady-state abundance profile. This method can be used in a predictive manner as well. Using aFUT to calculate the patients fractions of unique taxa pre-FMT to the donor sample and plotting that against the rrJSD for all post-FMT samples, the same trend as before holds; in other words, similar taxa implies a similar abundance profile, see Figure 6a4.

**Fig. 6.**
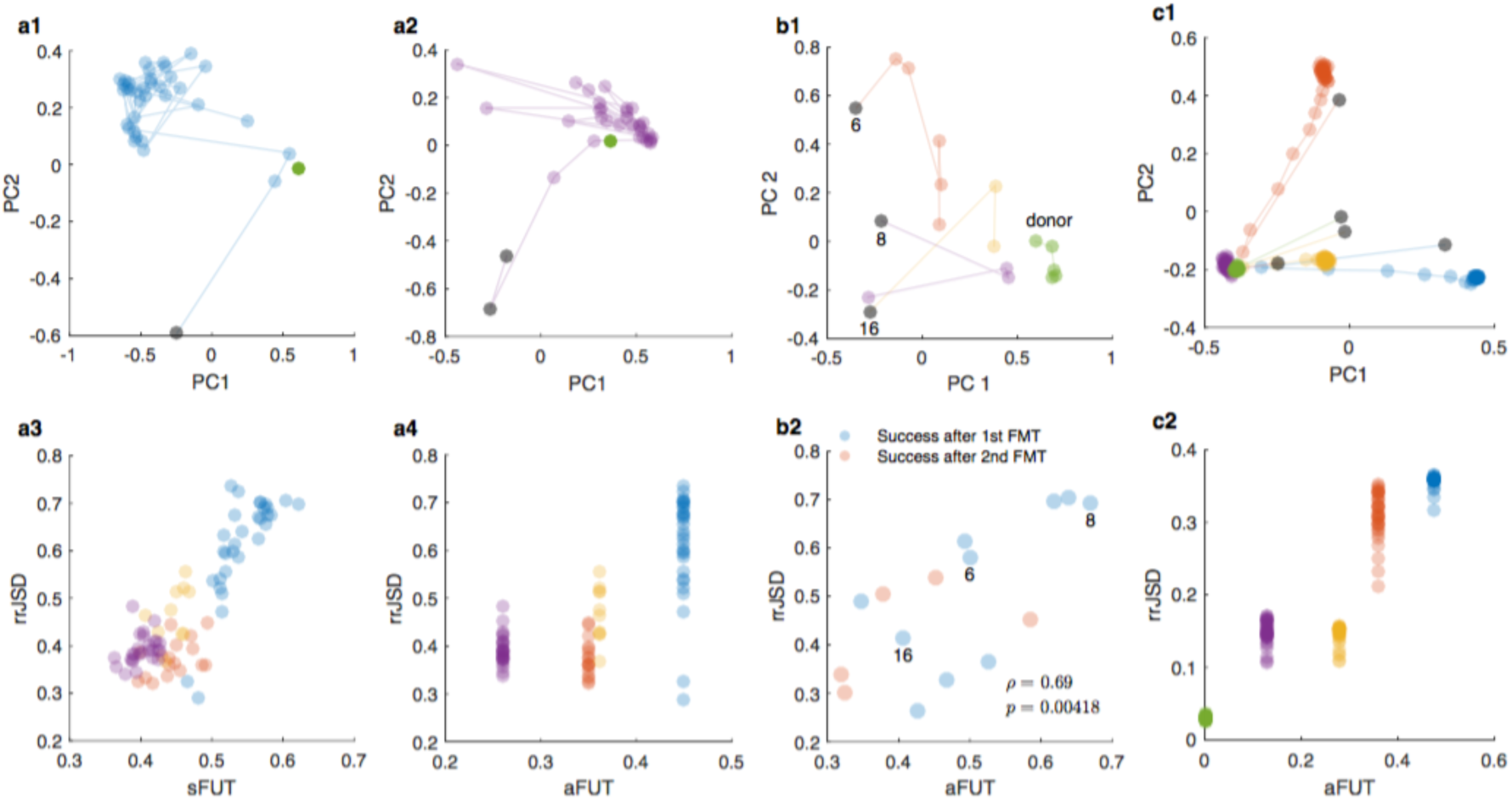
Perturbation by FMT (a) Long term study of four CDI patients receiving an FMT (3), n ∈ {35, 21, 14, 31} post FMT samples for the four respective patients {blue, orange, yellow purple}. (a1) Trajectory of patient 1, post-FMT shown in blue, pre-FMT shown in gray, and donor in green. (a2) Trajectory of patient 4, post-FMT shown in purple, pre-FMT shown in gray, and donor in green. (a3) Post-FMT samples paired with their four respective donor samples resulting in {35, 21, 14, 31} sample pairs. (a4) aFUT is computed from a single patient pre-FMT sample paired with the donor sample, and rrJSD is computed from post-FMT samples paired with donor samples. This is now predictive, in the sense that the x-axis only uses pre-FMT data and the y-axis is comparing post-FMT patient data to the donor. The first four time points post-FMT not included do to the fact that all patients are similar to the donor immediately after FMT, resulting in sample pairs of size {31, 17, 10, 27} (b) FMT study from MGH with n = 15 out of an original 20. (b1) Three representative trajectories for patients receiving an FMT. Patient 16 becomes most similar to the donor, patient 6 a little less similar, and patient 8, while temporarily being similar to the patient immediately post FMT does not remain similar (b2) aFUT is computed from a single pre-FMT sample paired with the donor sample, and rrJSD is computed from the last post-FMT sample, having been taken at least three days post FMT. Correlation coefficient and corresponding p-value displayed in the subfigure. Patients 6,8,16 highlighted. Correlation coefficient, ρ, and p-value (standard) displayed, see Methods section for details (c) Four synthetic patients shown in blue, orange, yellow and purple and a donor in green, made to mimic the Long term FMT study. (c1) Trajectories of all four patients, with the pre-FMT state shown in gray. (c2) aFUT is computed from a single pre-FMT sample paired with the donor sample, and rrJSD is computed from the post-FMT samples, excluding the samples corresponding to the transients immediately after FMT, see the Methods for more details on the in silico model.

The trend shown in Figure 6a was also observed in data from a larger study (36), where pre-FMT, donor, and post-FMT samples were available for 15 patients, see Figure 6b with the corresponding correlation coefficient and p-value shown in the subfigure. Patient 8 behaves in the exact same fashion as the patient highlighted in blue in Figure 6a. Both patients have a high aFUT with their respective donors, and they both immediately become similar to the donor immediately post-FMT, but then ultimately settle at a steady-state abundance profile that is far from the donor. Synthetic data using stochastic Generalized Lotka-Volterra (GLV) dynamics replicates the results as well, see Figure 6c.

## 4 Discussion

These results demonstrate that the human gut microbiome is an ecological system that has a preference toward a unique and asymptotically stable state for each collection of coexisting microbes. Identifying the asymptotic stability of the gut microbiome, in turn, allows us to make predictions about steady-state abundance profiles from the presence or absence of taxa alone. These findings hold promise as we move forward with personalized or precision microbiome-based therapies (37). One of the main themes reported in the literature is that our gut microbiome is highly individualized. Each person’s response to perturbations, such as antibiotics or FMT, is unique. We want to emphasize that this does not imply that we lack a complete understanding of the implications of that individuality. This paper illustrates that one of the major determinants of steady-state abundance is simply the microbes that are present. Therefore, given a potential donor pool, this work provides a first step towards choosing an optimal donor to minimize the dissimilarity between the patient’s post-FMT gut microbiome and the donor’s gut microbiome, (see Figure 1d for how our method could be applied). If we know that the post-FMT abundance profile of a patient can be better controlled by choosing a healthy donor for which the patient has the fewest possible unique taxa (low aFUT) then it is reasonable to assume that this will lead to a decreased probability of post-FMT complications. Indeed, if having less unique taxa compared to a donor leads to a more predictable post-FMT state, then one strategy for reducing sFUT between a patient and a donor would be to simply reduce the overall number of taxa in the patient by the administration of broad-spectrum antibiotics before FMT (24). This clean slate approach may hold promise for the general application of FMT in treating other complex diseases associated with a disrupted gut microbiome, such as, IBD, IBS and obesity.

## Acknowledgments

We acknowledge Simone Guglielmetti for sharing data with us. We also thank Aimee Milliken for very extensive editing. This work was partially supported by the John Templeton Foundation (award number 51977) and National Institutes of Health (R01 HL091528).

### Author contributions

Y.-Y.L. managed the project. Y-Y.L. and T.E.G. designed the project. T.E.G. developed the computational method, performed numerical simulations, and analyzed all the real data. T.E.G. and V.J.C. performed statistical tests. All authors analyzed the results. T.E.G., Y.-Y.L. and V.J.C. wrote the manuscript. All authors edited the manuscript.

